# Honey bees, *Apis mellifera*, are important pollinators of the highbush blueberry variety Ventura despite the inability to sonicate

**DOI:** 10.1101/551523

**Authors:** Keanu Martin, Bruce Anderson, Corneile Minnaar, Marinus de Jager

## Abstract

Animal-mediated pollination is an essential ecosystem service which over a third of the world’s agricultural crops depend on. Blueberry fruit production is highly dependent on pollinators and in their native range they are pollinated mostly by bumble bees (*Bombus* spp.). Demand for blueberries has increased in recent years due to their perceived health benefits. Consequently, blueberry cultivation has expanded well beyond their native range, including several regions where bumble bees are not present. In many areas, honey bees may be the only commercially available pollinators of blueberries because many countries ban the importation of bumble bees. This study aimed to determine the benefits of honey bee pollination on blueberry fruit quality and quantity for the variety Ventura by comparing yields of honey-bee-pollinated flowers to flowers where pollinators had been excluded. Honey bees significantly increased berry mass and diameter. Our results suggest that the presence of honey bee pollinators potentially increases revenue by approximately $864 501/ha in areas without bumble bees. We conclude that Ventura is reliably pollinated by honey bees, and that honey bee pollination may be a useful substitute for bumble bees in areas where bumble bees are absent. We also determined the extent to which blueberry yields could still be improved by comparing fruit quality and quantity under honey bee pollination to fruit quality and quantity achieved through ideal hand pollination. We found that blueberry yields may be still be significantly increased relative to ideal hand pollination and we discuss potential ways to improve the efficiency of honeybee pollination in the future. Additional research is required to study how beneficial honey bees are to fruit yield on varieties as the benefits of honey bees are likely to vary across different varieties.

## Introduction

Over 30% of the world’s agricultural crops depend on animal-mediated pollination, an essential ecosystem service which is valued at approximately €153 billion [1]. The dependence of crops on pollinators varies, with fruit formation in certain crops being extremely dependent on pollinators [2]. For example, atemoya, Brazil nut, cantaloupe, cocoa, kiwi, macadamia nut, passion fruit, pawpaw, rowanbarry, sapodilla, squashes and pumpkins, vanilla and watermelon, show a ca. 90% reduction in produce when pollinators are absent [2].

Blueberry production is also highly dependent on pollinators for the production of high-quality fruit [3–5]. This is a partly a result of floral architecture, where the pollen of blueberry flowers is concealed within poricidal anthers, making pollen transfer both within and between flowers unlikely without pollinators. For example, Campbell *et al*. [6] found that blueberry fruit were ca. 47% heavier when plants had access to pollinators (including honey bees, *Bombus* spp., *H. laboriosa* and *Xylocopa* spp.), than when pollinators were excluded. Effective pollinators of blueberries are typically large bees such as bumble bees [7–10], blueberry bees [6,11] and mining bees [7,8], which are able to extract pollen from anthers by vibrating their bodies at high frequency. This causes pollen to dehisce from the pores inside the blueberry anthers [11].

This “buzz pollination” strategy is employed by approximately 15,000–20,000 plant species [12,13]. Buzz pollination may have evolved to exclude less efficient or wasteful pollinators, such as honey bees (*Apis mellifera*), which are unable to obtain pollen from these specialized anthers through buzzing [7,10,12]. For example, Javorek *et al*. [7] found honey bees deposited approximately three times less pollen on blueberry stigmas during a single visit than bumble bees. However, the large numbers of foragers in honey bee colonies may enable effective pollination of blueberry flowers, as they may achieve increased flower visitation rates relative to bumble bees. This may explain why many commercial blueberry farmers completely, or partially, depend on honey bees as pollinators. Even within the native range of bumble bees, North American blueberry farmers frequently add honey bee hives to supplement bumble bee pollination [4,14–16].

The pervasive use of honey bees in the blueberry industry may also be a result of the prohibition on importation and use of non-native, commercially-produced bumble bees in parts of USA and many other regions where bumble bees are not native, such as southern Africa and Australasia [17,18]. These strict laws, prohibiting the movement of bumble bees, are in place because their introduction can have catastrophic effects on native fauna and flora [19,20]. For example, *Bombus ruderatus* and *Bombus terrestris* were introduced to Chile for agricultural pollination; these species have subsequently invaded southern South America, including Argentina, which has since banned commercial importation of bumble bees [21–24]. In Argentina, the highly invasive *Bombus terrestris* has caused a reduction in geographic range of the largest bumble bee in the world, and the sole native Patagonian bumble bee, *Bombus dahlbomii* [22]. Other potential impacts include pathogen transmission to native bumble bees, nectar robbing and flower damage [21,22,24]. Despite the potentially harmful effects of introducing bumble bees for agricultural pollination, the widely-held contention that honey bees are inferior pollinators of blueberries, drives the industry to place pressure on governments to allow bumble bee importation.

To alleviate the temptation to introduce bumble bees into new areas, it is pertinent to quantify the actual benefit of honey bees as commercial pollinators of blueberries, so that policies regarding the importation of bumble bees are based on substantive evidence and not impressions. Further, it may be possible to optimize the efficiency of honey bee pollination, so that the advantages of introducing bumble bees to new ranges are reduced. It is therefore important to quantify how well different blueberry varieties perform under honey bee pollination, while also estimating the potential for improvement by comparing blueberry yields under honeybee pollination to yields achieved under optimal hand pollination.

We aim to study the effects of honey bees on the production of blueberries in the variety Ventura, which is extensively planted in South America and South Africa [25]. More specifically, we compare the benefits (in terms of fruit quality, yield and revenue) of having honey bees as the only pollinators with blueberry yields achieved in the absence of pollinators. Despite their inability to buzz-pollinate, honey bees still transfer pollen between flowers and are capable of increasing fruit production in a variety of blueberry crops [4,5,26,27]. Consequently, we hypothesize that managed honey bees significantly increase fruit quality (i.e., mass and diameter) and decrease fruit development time, compared to flowers without access to pollinators. Next, we determine whether there is room to improve blueberry yields beyond that which is achieved when honey bee pollinators are allowed access to flowers. Although honey bees can transfer blueberry pollen between flowers [27], we expect that the inefficiency of honey bee pollination on blueberry flowers should result in significant potential for improvement, and consequently fruit quality and yield resulting from honey bee pollination should be lower than by hand pollination, which maximizes the transfer of pollen.

## Materials and Methods

This study was conducted in a one-hectare block of Ventura plants (7500) stocked with 15 honey bee hives on Backsberg blueberry farm (Western Cape, South Africa, 33°48’30.7“S 18°54’09.8”E). Our experiment consisted of three treatments: pollinator exclusion, open honey bee pollination, and optimized pollination (by hand). By comparing fruit quality among these three treatments, we determined whether the addition of honey bees was beneficial to blueberry production as well as the extent to which pollination by honey bees could potentially be improved upon (see Fig. 1 and treatment descriptions below). The three treatments were replicated across 20 plants, with each treatment applied once to each individual plant.

**Fig 1:**
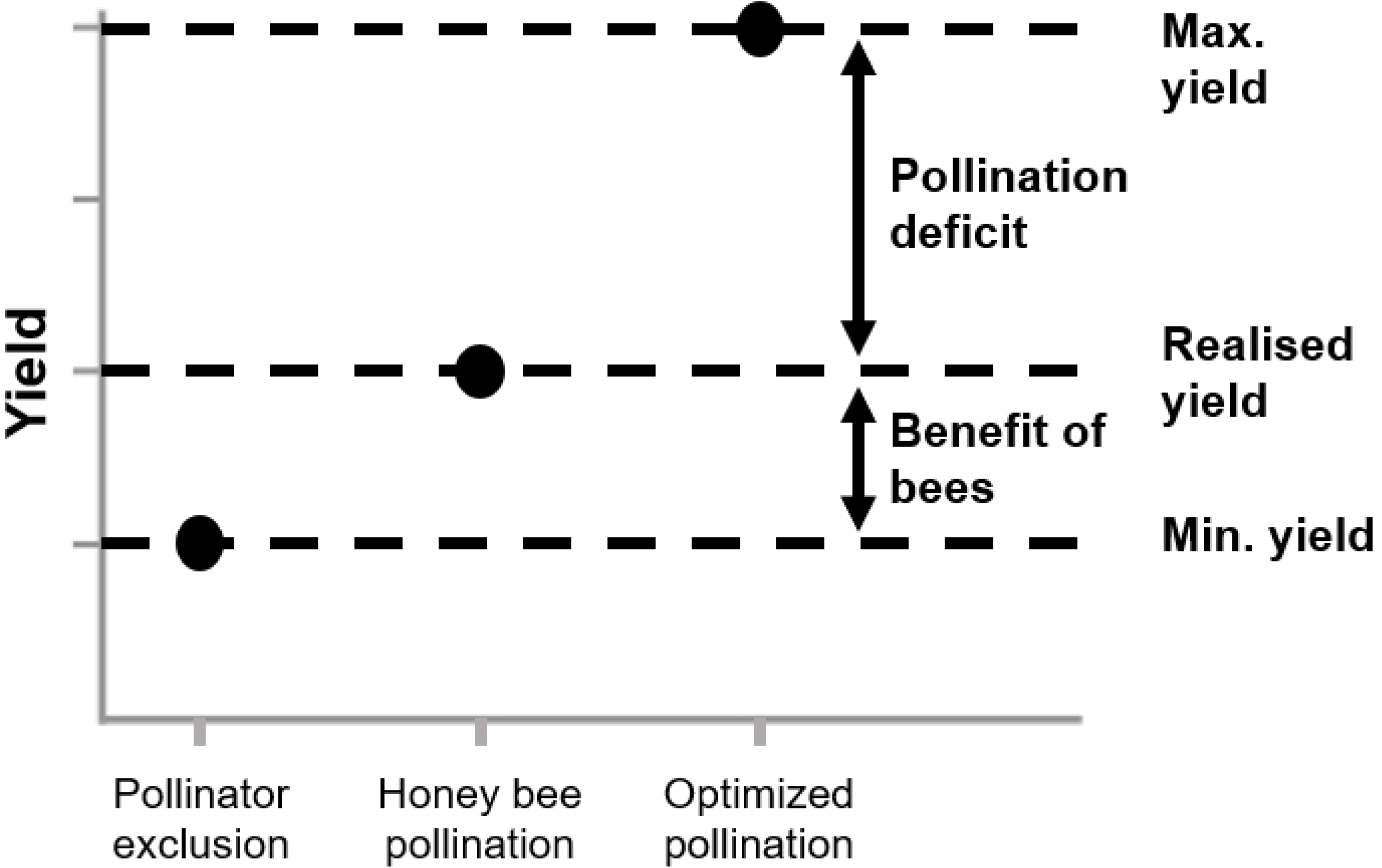
Hypothetical figure showing how the three treatments (pollinator exclusion, honey bee pollination and optimized pollination) may be useful metrics in determining the benefits of using honey bees to pollinate crops, and how yields could potentially be increased under more optimal pollination environments.

### Benefit of honey bees

To determine whether honey bees are commercially beneficial pollinators for berry production in Ventura, we compared blueberry production in plants where all pollinators were excluded (pollinator exclusion) with blueberry production in plants where honey bees had access to flowers (honey bee pollination) (See Fig. 1).

#### Pollinator exclusion (bagged)

Pollinators were excluded from visiting some blueberry flowers by placing a fine mesh bag over 20 individual virgin flowers (one flower per plant). Consequently, seed production in this treatment was the result of autonomous pollination and/or parthenocarpy (the production of fruit in the absence of fertilization), and not pollinator visitation. This provides an estimate of the yields expected in the absence of pollinators, and we expect that this should result in the poorest yield of our treatments, which we have termed “minimum yield”. Once flowers wilted, the bags were removed, and fruit maturation was allowed to continue normally.

#### Honey bee pollination (control)

Some flowers were left unbagged, allowing access to commercial honey bees placed on the farm. In South Africa, honey bees are generally the only pollinators observed on blueberries. The hive densities used for this treatment were 15 hives/ha, which corresponds to the densities of hives actually used by commercial blueberry farms [15,27,28]. Each flower was labelled to distinguish its fruit from other treatments. This treatment provides an estimate of the yield resulting from honey bee pollination and is expected to be similar to yields currently obtained by farmers of the Ventura variety across South Africa (realized yield).

### Pollination deficit

To determine the extent to which honey bee pollination can potentially be improved, we compared the honey bee pollination treatment (above) to a hand-pollinated treatment (optimized pollination). We assumed that hand pollination would result in the best fruit production possible by maximizing the deposition of pollen.

#### Optimized pollination (hand pollination)

Prior to hand-pollination, Ventura flowers were emasculated. We did this by removing part of the corolla with a scalpel, before removing all stamens with a pair of fine forceps, thus ensuring that no self-pollination could take place. To ensure that the experimental flowers were virgin, buds that were about to open were bagged three days before hand pollination. The day before pollination, pollen was collected from pollen donor flowers (approximately five flowers per pollen application) [3]. Pollen was extracted from donor flowers by removing the corolla with a scalpel and agitating the anthers with forceps, causing the poricidal anthers to release pollen into a Petri dish. Pollen from donor flowers on different individual plants was mixed together so that recipient flowers received pollen from multiple donors. This pollen mix was applied to recipient stigmas by dipping the stigma into the Petri dish containing pollen, and visually confirming that the stigma was saturated with pollen. Such careful hand pollination is likely to result in the maximum yield possible. Following hand pollination, a fine mesh bag was placed over hand-pollinated flowers to prevent honey bees from depositing additional pollen of unknown origin onto the stigma. This bag was again removed after the flower wilted.

### Measurements of fruit quality

After pollination treatments were applied, we checked fruit development once a week to determine whether fruit were mature and ready for fruit quality measurement. Fruits were considered mature when the entire fruit turned a uniform dark blue. Mature fruit were harvested by hand and subsequently weighed, and the diameter of each fruit was measured on the day it was harvested. By checking fruit every week, we could also determine the developmental period for each fruit (the number of weeks from pollination to harvesting) for each treatment. Apart from fruit mass and diameter, the developmental period is an important determinant of fruit quality, as early fruit are more valuable than late fruit; earlier fruit can be sold at higher prices when market demand is not saturated [3,29]. The percentage fruit set per treatment was also calculated.

### Estimating the economic impact of honey bees

In addition to fruit quality, we also determined how differences in pollination treatments could translate into differences in revenue gained. Firstly, to estimate the number of flowers, and thus potential fruits per plant, we used a constant flower number of 11 016 per plant [15]. This number serves only as an estimate of the total flowers for the highbush variety, Ventura, since we were unable to perform flower counts on our experimental plants. To calculate fruit yield per plant, we took the product of 11 016, the proportion of fruit set, and the predicted mass of fruits (taken from our linear mixed-effects model, see below) produced by individual plants for each treatment. Fruit yield allows us to determine whether differences between treatments found in fruit quality actually translates to yield, as it takes into account flowers that did not set fruit set as well. To calculate the per-hectare economic value of fruit for each treatment, we multiplied yield per plant by the number of plants per hectare and by the US dollars obtained per kilogram of fruit ($7.48) [30] (Eq. 1).

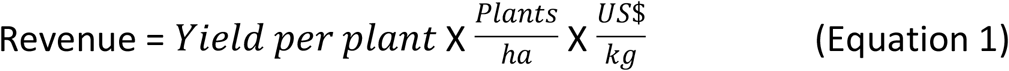

### Statistical analyses

To test for differences in fruit mass, diameter, and developmental period between treatments, we used a linear mixed-effects model with treatment nested in plant ID as the repeated random factor. Linear contrasts (Tukey) were used to test for treatment differences. To test how well our model explained the variance in our data, we used Nakagawa *R^2^* values [31], which provides both conditional variance (*R^2^c*) and marginal variance (*R^2^m*) estimates than can be equated to traditional *R^2^* values. Conditional *R^2^* values show the variance explained by the entire model (fixed effects and random effects), whereas marginal *R^2^* values show the variance explained by the random effects alone. The fixed effect was treatment and the random effect was treatment nested in plant ID. To test the overall effect of treatment on fruit set, we performed a log-likelihood ratio test between two mixed-effects logistic regression models, one with and one without treatment as a fixed effect. To test for differences in fruit yield (see calculation above) between treatments, we used a linear model. All statistical analyses were conducted in R (version 3.3.2) [32] using the packages nlme [33], multcomp [34], ggplot2 [35], sjPlot [36], car [37], lme4 [38], grid [32], gridExtra [39], lattice [40], MuMIn [41], plyr [42] and plotrix [43].

## Results

### Benefit of bees

The linear mixed-effects model explained the majority of the conditional variance for developmental period (*R^2^c*=0.999). The model also accounted for some of the marginal variance in developmental period (*R^2^m*=0.283), demonstrating that individual plants had different responses depending on treatment. The presence of honey bee pollinators did not significantly decrease the ripening period of blueberry fruits in comparison to treatments where honey bees were excluded (Table 1, Fig 2A).

**Fig 2:**
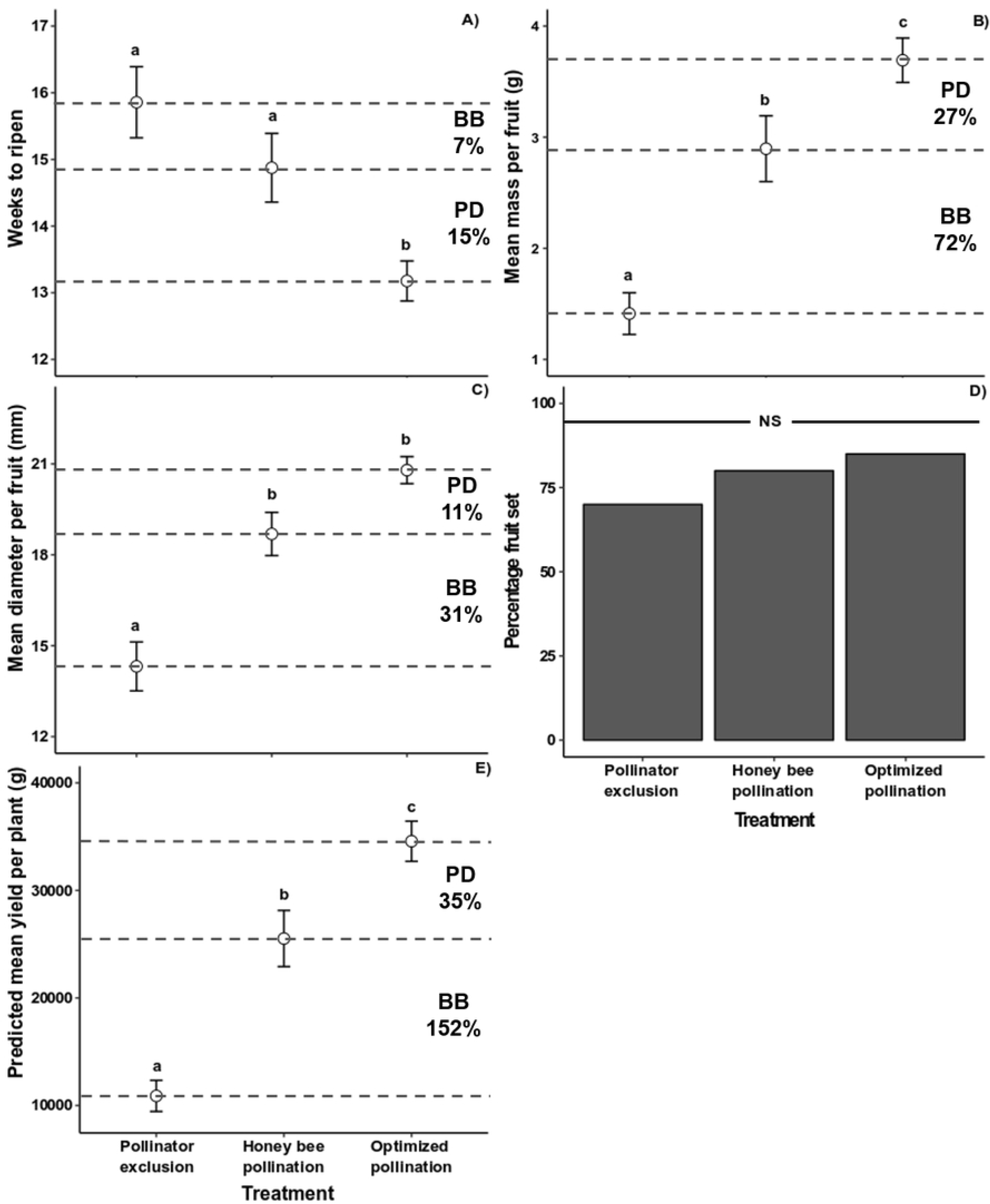
A) Mean number of weeks needed for the fruit to ripen for each treatment. B) The mean mass of fruits for each treatment. C) The mean diameter of fruits for every treatment. D) Percentage fruit set for each treatment. E) The mean yield per plant for each treatment. Letters indicate significance (*p*<0.05) of linear contrasts (Tukey HSD). Error bars indicate standard error. BB = benefit of bees, this is the percentage difference between flowers with no access to pollinators compared to flowers which had access to honey bees. PD = pollination deficit, this is the percentage difference between flowers with access to honey bees compared to flowers were hand pollinated.

**Table 1:** Outcome of three pollination treatments on blueberry fruit ripening, quality and yield.

|  | Benefit of bees (BB) | Pollination deficit (PD) | Pollinator exclusion – optimized pollination |
| --- | --- | --- | --- |
| Ripening period | $z=-1.505, p=0.2885$ | $z=-2.734, p=\mathbf{0.0173}$ | $z=-4.165, p<\mathbf{0.001}$ |
| Mass | $z=4.346, p<\mathbf{0.001}$ | $z=2.449, p=\mathbf{0.0381}$ | $z=6.771, p<\mathbf{0.001}$ |
| Diameter | $z=4.617, p<\mathbf{0.001}$ | $z=2.338, p=0.0508$ | $z=6.938, p<\mathbf{0.001}$ |
| Yield | $t=4.870, p<\mathbf{0.001}$ | $t=3.164, p=\mathbf{0.0078}$ | $t=7.992, p<\mathbf{0.001}$ |

Significance indicated by bold type

The linear mixed-effects model explained the majority of conditional variance for both mass (*R^2^c*=0.999) and diameter (*R^2^c*=0.999). The model explained a large proportion of the marginal variance for mass (*R^2^m*=0.503) and diameter (*R^2^m*=0.517), suggesting variation in individual plant responses in fruit mass and diameter per treatment. The presence of honey bees substantially increased fruit mass (mean ± SE fruit mass = 2.89g ± 0.23g, Table 1), with a 72% increase in mass per fruit compared to treatments without access to pollinators (mean ± SE fruit mass = 1.68g ± 0.25g, Fig 2B). Similarly, honey bees also caused a mean increase of 4mm (31%) per fruit (mean ± SE fruit diameter = 18.69mm ± 0.65mm) compared to treatments where honey bees were excluded (mean ± SE fruit diameter = 14.32mm ± 0.69mm, Fig 2C, Table 1). There was no difference in the fruit set of flowers with access to honey bees relative to flowers which had no access to honey bees or hand pollinated flowers (*Chi-square*=1.36, *df*=4, *p*=0.507) (Fig 2D).

The beneficial effects of honey bees on fruit mass and fruit set also translated into differences in total yield as calculated using Isaac’s [15] average flowers produced per plant, with the linear model explaining 54% of the variance in yield (R^2^=0.5355, F_(2,44)_ =27.52, p<0.001). Here, yield increased substantially (152%) from 10.11kg ± 2.2kg (mean ± SE predicted plant yield) per plant when pollinators were excluded to 25.52kg ± 3kg (mean ± SE predicted plant yield) when honey bees were allowed to forage on blueberries (Table 1, Fig 2E). Using a value of $7.48/kg [30] of blueberries and a density of 7500 plants per hectare, adding honey bee hives in areas lacking natural blueberry pollinators can potentially increase blueberry revenue by $864 501/ha (152%) compared to if pollinators were excluded.

### Pollination deficit

Hand pollination significantly shortened the ripening period of blueberry fruit by approximately two weeks or 15% (Table 1), from 15 ± 0.65 weeks (mean ± SE weeks to ripen) when pollinated by honey bees to 13 ± 0.64 (mean ± SE weeks to ripen) when pollinated by hand (Fig 2A). Hand-pollinated fruits were significantly heavier (Table 1) at 3.69g ± 0.23g (mean ± SE fruit mass), than flowers pollinated by honey bees at 2.89g ± 0.23g (mean ± SE fruit mass, Fig 2B), a ca.27% increase. Hand pollination did not significantly increase the size of fruits compared to fruits resulting from honey bee pollination (Table 1, Fig 2C).

However, when both mass and fruit set were incorporated into a model to calculate total yield, hand pollinations resulted in significantly greater yields than honey bee pollination (Table 1). Calculated per plant, optimizing pollination (i.e. hand pollination) can potentially increase yields from 25.52kg ± 3kg (mean ± SE predicted plant yield, after honey bee pollination) to 34.47kg ± 2.9kg (mean ± SE predicted plant yield, Fig. 2E), approximately 35%. This could lead to additional revenue amounting to $502 095/ha (35%).

## Discussion

This study revealed that the pollination environment has the potential to strongly affect the quality of fruit produced by the highbush blueberry variety Ventura. In particular, yields are greatly increased by the addition of honey bees in areas where bumble bee pollinators do not occur naturally, and importation is illegal. Honey bees were the only pollinators at this site and therefore the effects shown are a direct result of access to honey bees, rather than other unaccounted wild pollinators. This provides valuable information for the pollination of commercial blueberries, particularly with respect to the underutilized role played by honey bees, and suggests some important directions for research on blueberry pollination.

We show for the first time that Ventura can produce fruit without pollinators. However, these fruits are of lower quality than the fruits of flowers exposed to honey bee pollinators. This ability is not unique to Ventura, as other highbush blueberry varieties can also produce fruit in the absence of pollinators. However, these fruits are also of noticeably poorer quality than fruits produced by flowers with access to pollinators [5,6,15]. The presence of honey bees significantly increased blueberry yields by improving fruit quality through greater fruit diameter and mass (Fig. 2). This shows that despite honey bees’ inability to buzz-pollinate, they do extract blueberry pollen from anthers and transfer it to stigmas. Thus, in areas lacking native blueberry pollinators, the addition of honey bees may increase blueberry yields by more than 150% (Fig 2E). This translates to an economic value of approximately $864 501/ha. Consequently, honey bees may be extremely beneficial, potentially eliminating the need to import bumble bees in countries which do not have native blueberry pollinators.

Despite this benefit, there is still a pollination deficit of approximately 27%, which suggests that there may be room to optimize pollination. However, it is unclear exactly why honeybee pollination results in sub-optimal fruit yields and whether native bumble bee pollinators result in greater yields than high densities of honey bees. These represent important directions for future research on blueberry pollination. There could be several reasons for the sub-optimal yields produced by honey bee pollination and future research needs to concentrate on these to optimize the pollination environment. Below we discuss four potential reasons for sub-optimal yields, each of which should be targeted in future studies in an attempt to improve blueberry yields.

### Hand pollinations are not a realistic maximum yield

The magnitude of the pollination deficit is contingent on what honey bee-pollination yields are being compared with (in this case hand-pollinations). It is possible that blueberry yields resulting from honey bee pollination are already close to the maximum that can be achieved through animal pollination and that no amount of tinkering is likely to reduce this perceived deficit. For example, pollination by bumble bees may not result in smaller pollination deficits than pollination by honey bees. Unfortunately, there is a lack of data comparing blueberry fruit yields after pollination by different bee species to fruit yields after hand pollinations. Consequently, it is unclear how pollination deficits under different pollinator environments are likely to vary and if honey bee hives are any less effective than bumble bee colonies as commercial pollinators of blueberries. This represents an important first step in determining whether the pollination deficit can be reduced.

### Blueberry attractiveness

If honey bee pollination really is less effective than other modes of pollination, it may be the result of low visitation rates to blueberry flowers. Blueberry flowers may be less attractive to honeybees than wild or other agricultural flowers that surround blueberry farms. Low attractiveness relative to other flowers may occur because blueberries are adapted to larger bees and both their nectar and pollen rewards may be more difficult for honeybees to access [44]. Ventura has a long floral tube length (11.39mm ± 0.4mm) which may make it difficult for honey bees to access nectar at the bottom of the flower as bees would need to insert nearly half of their bodies into the flower to reach the nectar. This increases honey bees energy expenditure and may cause honey bees to search for more favourable flowers. A possible way to overcome this is through the development of varieties with shorter or wider corollas.

### Honeybees may not pick up pollen

High visitation by honey bees may still result in low fruit set if they forage for nectar but remove very little pollen from the poricidal anthers. This could occur because honey bees are not capable of buzz pollinating and hence extract few pollen grains per visit [7,10,12]; honey bees do appear to deposit very few pollen grains per visit to blueberry flowers [7]. Other than trying to develop varieties with pollen which is more easily released, there is very little one can do to improve this potential problem.

Further research needs to explore the efficiency of honey bees as commercial pollinators of different blueberry varieties, especially comparative studies with bumble bees. More detailed investigations of behavioural interactions of honey bees with blueberry flowers may highlight traits which determine honey bee preferences for different varieties. Studies on the mechanical fit of honey bees to the flowers of different blueberry varieties may also illuminate which varieties are best suited to honey bee pollination.

## Acknowledgements

We would like to thank BerryWorld South Africa for providing access to the Backsberg farm. We would also like to thank our funders, the National Research Foundation (South Africa) under grant number 112277 and South African Berry Producers Association (KM), the Claude Leon Foundation (MDJ), the Eva Crane Trust (ECTA_20170609 to CM and ECTA_20170905 to MDJ) and the National Research Foundation (South Africa) (105987 to BA and 111979 to CM).

